# High-efficiency transformation and expression of genomic libraries in yeast

**DOI:** 10.1101/2023.08.07.552314

**Authors:** Mira Loock, Luiza Berenguer Antunes, Rhiannon T Heslop, Antonio Alfonso De Lauri, Andressa Brito Lira, Igor Cestari

## Abstract

*Saccharomyces cerevisiae* is a powerful system for the expression of genome-wide or combinatorial libraries for diverse types of screening. However, expressing large libraries in yeast requires high-efficiency transformation and controlled expression. Transformation of yeast using electroporation methods is more efficient than chemical methods; however, protocols described for electroporation require large amounts of linearized plasmid DNA and often yield about 10^6^ cfu/µg of plasmid DNA. We optimized the electroporation of yeast cells for the expression of whole-genome libraries to yield up to 10^8^ cfu/µg plasmid DNA. The protocol generates sufficient transformants for 10-100x coverage of diverse genome libraries with small amounts of genomic libraries (0.1µg of DNA per reaction) and provides guidance on calculations to estimate library size coverage and transformation efficiency. It describes the preparation of electrocompetent yeast cells with lithium acetate and dithiothreitol conditioning step and the transformation of cells by electroporation with carrier DNA. We validated the protocol using three yeast surface display libraries and demonstrated using nanopore sequencing that libraries’ size and diversity are preserved. Moreover, expression analysis confirmed library functionality and the method’s efficacy. Hence, this protocol yields a sufficient representation of the genome of interest for downstream screening purposes while limiting the amount of the genomic library required.

## 1. Introduction

The transformation of *Saccharomyces cerevisiae* is a widely used method of genetic modification whereby exogenous DNA is inserted into a cell. Yeast cells can express proteins in their native folds and undergo post-translational modifications, offering further benefits over other bacterial recombinant methods [1]. Intact cells have been transformed using various methods, including glass beads, lithium, and electroporation [2]. Despite there being several protocols available for the transformation of yeast cells, most approaches are not suitable for transforming yeast with large genome-wide or combinatorial libraries, which often contain 10^6^ to 10^9^ plasmids or DNA fragments [3]. Transforming yeast cells with large libraries requires a high-efficiency transformation with an order of magnitude equivalent to or higher than a typical library size. Previously published high-efficiency methods transform yeast with linearized vectors or PCR products containing the sequence coding the peptide of interest [3]. A high-efficiency method was described to assemble libraries in yeast [3]. Although efficient, it has many drawbacks, such as requiring a large quantity of DNA, ∼12µg, and problems for downstream protein isolation as multiple fragments can recombine in a single yeast cell [1,3].

We aimed to generate a protocol for efficient transformation of assembled genome-wide or combinatorial libraries for expression in yeast. We used libraries to express genomic fragments of parasitic pathogens – *Trypanosoma brucei, Trypanosoma cruzi*, and *Giardia lamblia* for yeast surface display [4], but the protocol can be used for the transformation of any library or DNA vector requiring high-efficiency transformation. In our approach, libraries were cloned in pYD1 vector for galactose induction and surface expression in *S. cerevisiae* EBY100 strain using the Aga1p-Aga2p protein display system [5]. The plasmid DNAs were transformed in yeast by electroporation, which forms transient pores in the cell membrane for efficient DNA uptake [6]. We optimized DNA intake using DNA carriers with nanograms of plasmid DNA and describe methods for long-term storage and induction of library using a galactose-inducible system. We demonstrate the efficacy of the method to express three different genome-wide libraries. We sequenced the libraries before and after yeast transformation by Oxford nanopore sequencing and demonstrated that the method preserved library size and diversity. Moreover, Western blot and flow cytometry analysis confirmed library expression in yeast. Hence, the protocol yields a transformation efficiency of up to 10^8^ cfu/µg DNA, yet only requires 0.1µg of DNA per reaction, thus reducing the amount of DNA and transformation reactions required for whole genomic library representation.

## 2. Experimental Design

Before beginning the experiment, plasmid DNA was obtained using Mini-Prep kits and further cleaned up using size-selection magnetic beads. The cleanup results in DNA with purity of about 1.8 260/280 and 2.0 260/230 (see note 1). We optimized this protocol by using 100 ng of DNA of libraries constructed in the pYD1 vector to minimize library usage. This system uses the galactose-inducible GAL1 promoter to induce the expression of surface proteins. Transformants are selected by growing on synthetic-defined (SD) media lacking tryptophan. This allows for the selection of successfully transformed yeast with the TRP1 selectable marker.

We recommend electroporating yeast without plasmid and growing on YPD to ensure that yeast can survive treatment with buffers or electroporation. We also recommend growing yeast electroporated without plasmid on SD/-trp or respective selection marker to ensure that there is no contamination or unspecific growth associated with the yeast selection system. To evaluate transformation efficiency, yeast should be electroporated with a known plasmid (ex. pYD1) and grown on SD/-trp. If all controls, as seen in Table 1, give the desired outcome, we recommend transforming with genomic libraries or DNA of interest.

**Table 1.** Experimental conditions and its expected outcomes for the evaluation of transformation efficiencies. Dex, dextrose.

| Experimental condition | Expected Outcome |
| --- | --- |
| Yeast plated on YPD agar | Growth |
| Yeast plated on SD/-trp dex agar | No growth |
| pYD1 plated on SD/-trp dex agar | Growth |
| pYD1 + genomic library plated on SD/-trp dex agar | Growth |

### 2.1. Materials

#### 2.1.1. Preparation of plasmid DNA

1. Nucleospin plasmid mini-prep kit (Takara Bio USA Inc., catalogue number 740588.50)
2. NGS clean-up and size selection magnetic beads (Takara Bio USA Inc., catalogue number 744970.5)

#### 2.1.2. Preparation of Electrocompetent cells

1. 1000µl pre-sterile barrier tips (Neptune, catalogue number BT100.96)
2. Saccharomyces cerevisiae Meyen ex E.C. Hansen, EBY100 strain (ATCC, catalogue number MYA-4941)
3. Yeast extract (Bio Basic, catalogue number G0961)
4. Peptone (Bioshop Canada Inc., catalogue number PEP403.500)
5. Dextrose anhydrous (Fisher Scientific, catalogue number D16-500)
6. D-Sorbitol (Bio Basic, catalogue number SB0491.SIZE.500g)
7. Calcium chloride (Fisher Scientific, Acros Organics, catalogue number 349610025)
8. Lithium acetate (Fisher Scientific, Acros Organics, catalogue number AC297111000)
9. Dithiothreitol (DTT) (Bio basic, catalogue number DB0058.SIZE.5g)

#### 2.1.3. Transformation of electrocompetent cells

10. 1000µl pre-sterile barrier tips (Neptune, catalogue number BT100.96)
11. 200µl pre-sterile barrier tips (Neptune, catalogue number BT200)
12. 10µl pre-sterile barrier tips (Neptune, catalogue number BT10)
13. pYD1 vector (Addgene, catalogue number 73447)
14. Agar A (Bio Basic, catalogue number FB0010)
15. Cuvettes plus 2mm gap (Fisher Scientific, catalog number FB102)
16. Semi-micro cuvette, PS (Sigma-Aldrich, catalogue number BR759015-100EA)
17. Falcon 50mL Conical Centrifuge Tubes (Fisher Scientific, catalogue number 14-432-22)
18. 100mm x 20mm Tissue Culture Treated Dishes (Ultident Scientific, catalogue number 229621)

#### 2.1.4. Freezing transformed cells

1. Glycerol (Fisher Scientific, catalogue number BP229-1)
2. 2ml Microcentrifuge tubes with screw caps (Fisher Scientific, catalogue number 02-682-558)

#### 2.1.5. Induction of YSD

1. Yeast nitrogen base w/o AAs, w/o ammonium sulphate (Bioshop Canada Inc., catalogue number YNB404.250)
2. DO supplement -trp (Takara Bio USA Inc., catalogue number 630413)
3. Ammonium sulphate (Fisher Scientific, catalogue number A702-500)
4. D-(+)-raffinose pentahydrate (Fisher Scientific, catalogue number R000225G)
5. D-Galactose (Fisher Scientific, catalogue number BP656500)

### 2.2. Equipment

- Bunsen burner (Fisher Scientific, catalogue number M-650713-2463)
- AB15 Accumet® basic pH meter (Fisher Scientific, catalogue number 11376202)
- Avanti® J-E BioSafe High Performance Centrifuge (Beckman Coulter, catalogue number A20699)
- Stackable incubator shaker (New Brunswick™, catalogue number M1282-0012)
- Visible spectrophotometer V1-200 (VWR, catalogue number 634-6000)
- Gene Pulser Xcell Microbial System (Bio-Rad, catalogue number 1652662)
- Vortex mixer (Fisher Scientific, catalogue number 02215365)
- Freezer -80°C (New Brunswick™, catalogue number U9430-0001)

## 3. Procedure

### 1. DAY ONE

#### A) Calculate the number of required library transfections

1. Calculate the number of required transfections based on the required number of transformants, i.e., colony forming units (cfu), to obtain 10 to 100-fold the library size. If the library size is 10^6^ clones, then 10^7^ (10-fold) to 10^8^ (100-fold) yeast transformants are required. In our experience, aiming for 10-fold, or more, for the library size increases the library representation in yeast (See Fig 2, and Expected Results). Use the formula below to calculate the number of electroporations required to generate the number of required yeast transformants.

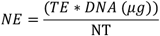

Where NE is the number of electroporations, TE is the transformation efficiency, and NT is the required number of transformants. We recommend performing a few transformations to verify transformation efficiency. It should be between 10^7^ - 10^8^ cfu/µg DNA using this protocol.

**Figure 1.**
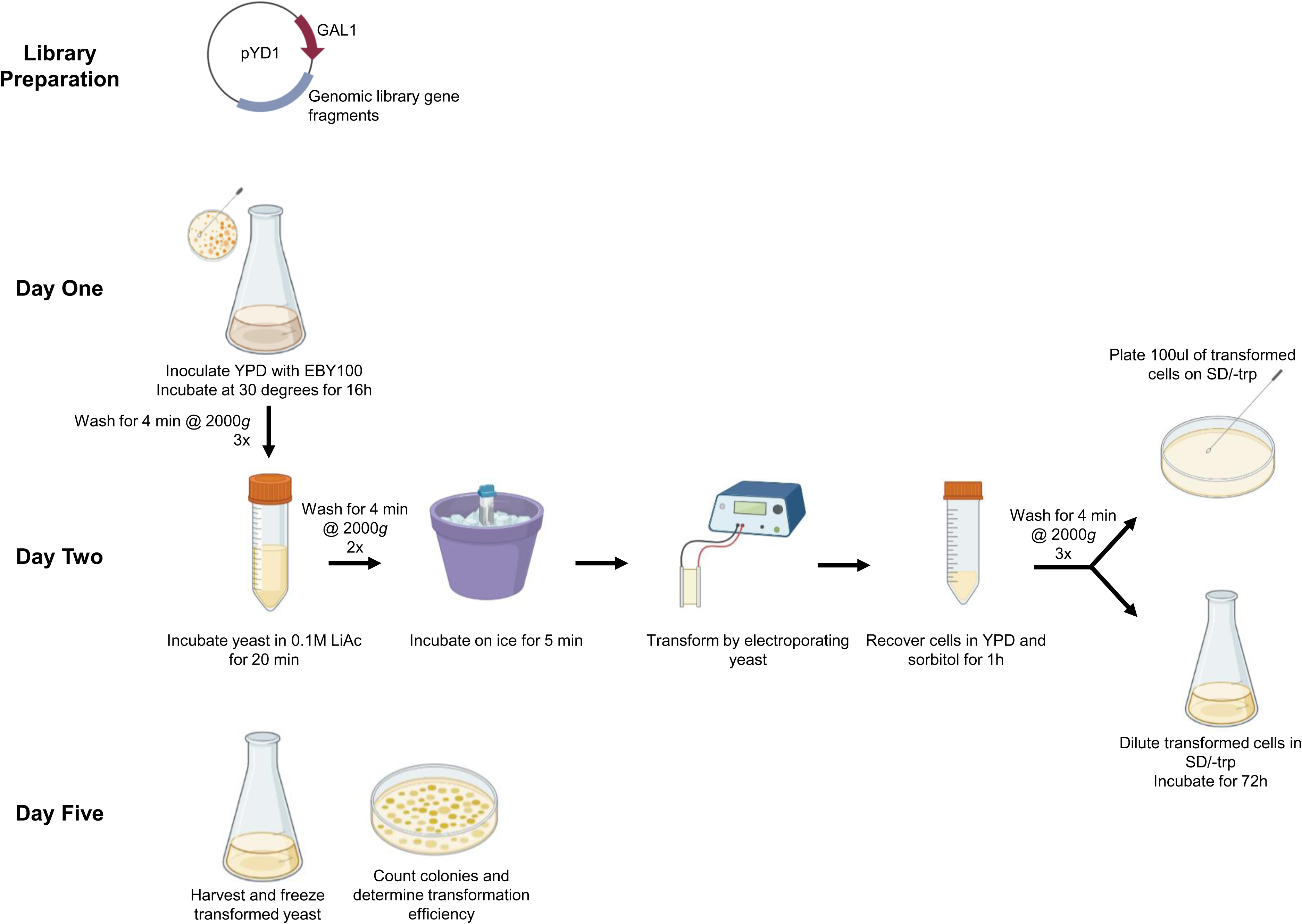
Overview of experimental design. Prepare genomic library in plasmid vector. On day one, inoculate and grow yeast. Day two, condition the yeast in 0.1M Lithium Acetate (LiAc) and electroporate. Recover cells in YPD medium and select transformants by plating dilutions in SD/-trp dex agar plate (top) and expanding transformed yeast in SD/-trp dex culture (bottom). Day five, harvest the transformed yeast for freezing or experiments, and count the colonies on agar plates to calculate transformation efficiency.

**Figure 2.**
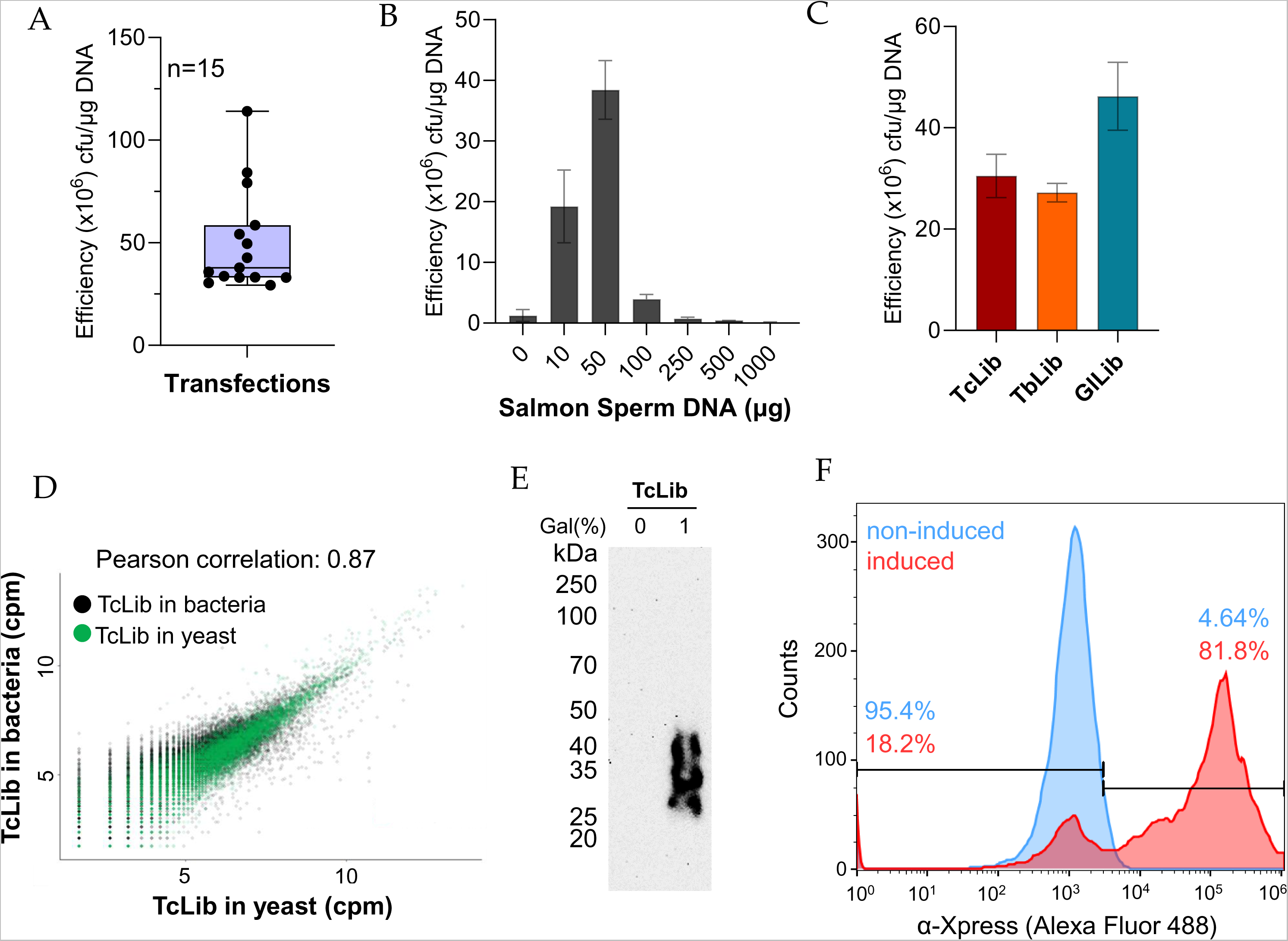
Expected results. A) Range of transformation efficiency of yeast EBY100 strain using pYD1 or genomic libraries constructed in pYD1. Number of experiments (n) is 15. B) Effect of salmon sperm DNA on transformation efficiency. C) Transformation efficiency obtained with three genome-wide libraries, TcLib, *T. cruzi* library; TbLib, *T. brucei* library; GlLib, *G. lamblia* library. B-C, data show the mean of at least three biological replicates ± standard deviation. D) Correlation of library sequencing after DNA extraction from bacteria (used to generate libraries) or yeast (used to express the libraries). E) Western blot with monoclonal mouse α-Xpress (1:1,000 dilution, Life Technologies) antibodies against Xpress tag fused to library proteins. Western was developed by chemiluminescence after incubation with goat α-mouse IgG-HRP. F) Flow cytometry analysis with non-induced (glucose) and induced (1% galactose) library expression in yeast. Cells were labelled with monoclonal mouse α-Xpress antibodies (1:500 dilution) and goat α-mouse IgG-Alexa Fluor 488 (1:1,000 dilution, Life Technologies).

#### B) Prepare solutions (see topic 5, Reagents Setup) and *S. cerevisiease* EBY100 strain culture

1. Prepare and sterilise all solutions. 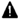**CRITICAL STEP** The 0.1M LiAc/10mM DTT filter sterilized solution should be made fresh on the day of electroporation.
2. Prepare at least 4 SD/-trp dex agar plates per library transfection.
3. Take 1 colony (∼2 mm) of *S. cerevisiease* EBY100 strain from working stock plate (see notes 2 and 3) and inoculate into 5ml of sterile YPD. Allow to grow for approximately 7 hours in an incubator shaker at 30°C whilst shaking at 225rpm.
4. Add the 5ml culture to 95ml fresh YPD to reach an OD600 of 0.02-0.07 and allow to grow overnight (16h) at 30°C whilst shaking at 225rpm (see note 4).

### 2. DAY TWO

#### A) Preparation of electrocompetent cells

1. Prepare 100ml EBY100 culture in YPD per 4 electroporation reactions. Measure the OD600 of the overnight culture, the optimal OD600 range is between 1.6-1.8 after 16 hours in a 30°C incubator whilst shaking at 225rpm. This will depend on the doubling time of your yeast (see note 5).
2. Collect EBY100 cells by centrifugation at 2000g for 4 mins, at 4°C and remove the media.
3. Ressuspend by vortexing the cell pellet in 50ml sterile ice-cold ddH2O per 100ml of culture. Centrifuge at 2000g for 4 mins, 4°C. Repeat this step once.
4. Ressuspend by vortexing the cell pellet once in 50ml ice cold electroporation buffer per 100ml of culture. Centrifuge at 2000g for 4 mins, 4°C.
5. Resuspend by vortexing the cell pellet in 20ml of fresh, 0.1M LiAc/10mM DTT (pre-warmed to 30 degrees) per 100ml of culture. Transfer the cell suspension to a covered Erlenmeyer flask and condition the yeast by shaking for 20 mins at 30°C, 225rpm (see note 6).
6. Collect cells by centrifugation at 2000g for 4 mins, 4°C.
7. Ressuspend by vortexing the cell pellet in 50ml ice-cold electroporation buffer per 100ml of culture. Centrifuge at 2000g for 4 mins, 4°C.
8. Resuspend by vortexing the cell pellet in ice-cold electroporation buffer to achieve a final volume of 1.4ml per 100ml of culture. Keep the cells on ice until electroporation. There should be approximately 1.5 × 10^9^ cells, which is sufficient for 4 electroporation reactions of 200µl each (2.1×10^8^ cells per reaction).

#### B) Transformation of electrocompetent cells with genomic library

1. Prepare 5ml of 1:1 Sorbitol:YPD per electroporation and store on ice.
2. For each electroporation prepare 0.1µg of genomic library DNA, 25µg of Salmon Sperm DNA and 200µl of electrocompetent cells, mix gently by pipetting and transfer to a pre-chilled 2mm cuvette. Prepare a separate negative transformation control using 0.1µg of pYD1 plasmid per 200µl electrocompetent cells (see note 7).
3. Incubate the cells on ice for 5 mins.
4. Electroporate cells at 2.5kV, 25µF, 200Ω with a Gene Pulser Xcell Microbial System. The time constant ranges should be between 3.1-4.2 ms.
5. Immediately add 1ml of pre-chilled Sorbitol:YPD to the cuvette. Transfer the cells to a 50ml centrifuge tube containing 3ml of Sorbitol:YPD. An additional flush of the cuvette with another 1ml of sorbitol:YPD will help recover cells from the cuvette. Adjust the final volume to 5ml Sorbitol:YPD solution. 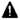 **CRITICAL STEP** Keep the cells on ice until all the electroporations are finished and cells are resuspended in Sorbitol:YPD.
6. Allow transformed cells to recover for 1 hour by placing the tubes in an incubator shaker at 30°C whilst shaking at 225rpm.
7. Harvest cells by centrifugation at 2000g for 4 mins, at 4°C.
8. Resuspend by vortexing the cell pellet in 10ml ddH2O to eliminate any trace of YPD. Centrifuge at 2000g for 4 mins, 4°C. Repeat this step twice.
9. Resuspend by vortexing cells in 4ml SD/-trp dex.
10. **OPTIONAL STEP** Prepare serial dilutions of two samples of each library transfection in SD/-trp dex and plate 100µl of 1:50 and 1:100 dilutions on SD/-trp dex agar plates for calculation of transformation efficiency. Invert plates and place them in 30°C incubator for 3-5 days, until colonies are visible.
11. The cultures transformed with the same genomic library can be pooled together and resuspended in a 1:30 dilution of SD/-trp dex media to an OD600 around 0.1-0.2. A volume of roughly 150ml is typically used as a 1:30 dilution that reaches the expected OD600.
12. Allow the culture to expand for 48 hours by incubating on a platform shaker at 30°C whilst shaking at 225rpm.
13. Check the OD600 of the culture to ensure the culture has expanded (see note 8). The OD600 is expected to be around 1.5-1.8. A;t this point, transformed cells can be frozen or surface protein expression can be induced.

#### C) Freezing transformed yeast cells

1. Harvest cells by centrifugation at 2000g for 4 mins at 4°C.
2. Resuspend by vortexing pellet in 1ml of SD/-trp dex with 25% glycerol per 10ml of culture. The glycerol causes the cells to freeze slowly reducing damage caused by the formation of ice crystals.
3. Freeze cells in a -80°C freezer.

### 3. DAY FIVE

#### A) Calculate the transformation efficiency and genome coverage of the YSD library

1. Check the SD/-trp dex agar plates for visible colonies and count the total number of colonies on the plates. Count plates from dilutions resulting in 30 to 300 colonies.
2. Calculate the transformation efficiency using the following formula [9]:

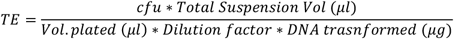
3. Typical transformation efficiencies range between 10^7^ to 10^8^.

#### B) Thawing transformed yeast cells

1. Thaw each vial of transformed yeast in 37°C water bath for 30 seconds.
2. Take 200ul of cells, ∼6×10^6^ cells and recover in 5ml YPD at 30°C and 225rpm for 2 hours or until an OD600 of 1.
3. Harvest cells by centrifugation at 2000g for 4 mins at 4°C.
4. Ressuspend by vortexing cells in 10ml ddH2O and centrifuge at 2000g for 4 mins at 4°C to remove any trace of YPD. Repeat this step twice.
5. Resuspend by vortexing cells in 20ml selective media (SD/-trp dex) to an OD600 of 0.4 and allow the culture to grow for 4 hours or until an OD600 of ∼1 in an incubator shaker at 30°C shaking at 225rpm.

#### C) Induction of YSD

1. Harvest cells by centrifugation at 2000g for 4 mins at 4°C.
2. Ressuspend by vortexing cells in 10ml ddH2O and centrifuge at 2000g for 4 mins at 4°C. Repeat this step once.
3. Resuspend by vortexing the cell pellet in 20ml SD/-trp/raf media. Grow at 225rpm, 30°C for 2 hours to adapt cells to raffinose.
4. Harvest cells by centrifugation at 2000g for 4 mins at 4°C.
5. Resuspend by vortexing pellet in 20ml of SD/-trp/raf/ 3% galactose. Incubate for at least 16 hours at 225rpm 30°C to induce the expression of the library (see note 9).
6. **OPTIONAL STEP** To validate protein expression by Western blot, take an aliquot of induced cells, lyse the cells and extract proteins according to protocol [7]. Perform western blot using antibodies against the Xpress and V5 tag, or against proteins expressed in the library [8].

### 3.1. Notes

1. The quality of the DNA used can greatly affect transformation efficiencies, thus, cleaning up with magnetic beads is highly recommended.
2. All steps manipulating the yeast should be carried out in sterile conditions, such as in the vicinity of a flame.
3. Always use an EBY100 colony from a fresh (<4 weeks old) working stock plate. Avoid selecting too small or too large colonies. If a fresh plate is unavailable, streak a new one from freezer stock EBY100 and not from a previous working stock plate.
4. Yeast will grow faster with better aeration. It is recommended to shake them at 220-270rpm whilst growing.
5. We strongly recommend performing a growth curve of your EBY100 to establish the doubling time of EBY100 in the conditions specific to the user’s laboratory.
6. We have found that conditioning the cells in LiAc/DTT for longer than 30 minutes significantly decreases transformation efficiency.
7. Changing the amount of DNA in electroporation can affect transformation efficiency.
8. At this transformation efficiency, approximately 99.5% of the cells die, which typically shows as clumps of black debris in the culture. It does not affect the procedure.
9. Yeast galactose induction requires media with no glucose. In this example, the Aga1p-Aga2p display system using pYD1 and EBY100 yeast strain uses a GAL1-activated promoter, which is repressed by the presence of glucose and induced by galactose. 2% raffinose is used as an alternative carbon source in the SD/-trp dex media to avoid affecting GAL1 induction.
10. To avoid diluting the media by adding galactose prepared in water, consider preparing 30% galactose stock in SD/-trp dex media. D-galactose solubility in water is 32%.

## 4. Expected Results

Our transformation protocol gave a transformation efficiency of 10^7^ to 10^8^ transformants/µg of library DNA with the average transformation efficiency being 4.6×10^7^ transformants/µg of library as seen in Figure 2A. The use of carrier DNA has been shown to improve transformation efficiency [9]. We determined that 50µg of salmon sperm DNA consistently increased the efficiency to ∼5×10^7^ transformants/µg DNA which can be seen in Figure 2B. The amount of salmon sperm DNA may need to be optimized for variations between batch or suppliers, as we noted variations between products from different suppliers. We tested the transformation protocol with three different genomic libraries from protozoan parasites – *Trypanosoma cruzi, Trypanosoma brucei*, and *Giardia lamblia*. The average transformation efficiency was 3.05×10^7^, 2.72×10^7^, and 4.62×10^7^ transformants/µg DNA, respectively, as seen in Figure 2C. This indicates reproducibility amongst genomic libraries independent of DNA origin. We sequenced the library DNA extracted from DH5α *E. coli* (used to generate the library) and the genomic library extracted from *S. cerevisiae* (used to express the library) using Oxford Nanopore Sequencing to confirm the complete transformation of the library size. We found a Pearson correlation of 0.866, suggesting maintenance of library size as seen in Figure 2D, and the sequence analysis showed that library diversity was also preserved [10].

In Figure 2E, protein expression of yeast transformed with the genome-wide *T. cruzi* library (TcLib) was validated by western blot. A smear is seen when yeast are grown in the presence of galactose inducing protein expression. This is not seen in the absence of galactose. In Figure 2F, to further validate the presence of surface proteins, flow cytometry was conducted on the transformed TcLib. When yeast are grown in the presence of galactose, 81.8% of the cell population expressed the α-Xpress tag on the surface of the yeast, indicating the presence of surface proteins.

## 5. Reagents Setup

1. YPD
  a. Dissolve 10g yeast extract and 20g peptone in 850ml MilliQ H20.
  b. Adjust pH to 6.5 with 1M HCl or 1M NaOH.
  c. Add MillQ H_2_0 to final volume of 900ml.
  d. Autoclave for 20min at 121°C.
  e. Wait to cool, then add 100ml of sterile 20% dextrose under sterile conditions (e.g., under a flame) and store at 4°C.
2. SD/-trp dex
  a. Dissolve 1.7g yeast nitrogen base, 5g ammonium sulphate, 0.74g -trp drop out supplement in 850ml MillQ H_2_O.
  b. Adjust pH to 5.8.
  c. Add MillQ H_2_O to final volume of 900mL.
  d. Autoclave for 20min at 121°C.
  e. Wait to cool, then add 100ml of sterile 20% dextrose under sterile conditions and store at 4°C.
3. SD/-trp dex agar
  a. Dissolve 1.7g yeast nitrogen base, 5g ammonium sulphate, 0.74g -trp drop out supplement in 850ml MillQ H2O.
  b. Adjust pH to 5.8.
  c. Add 20g agar.
  d. Add MillQ H_2_O to final volume of 1L.
  e. Autoclave for 20min at 121°C and store at room temperature.
  f. Wait to cool, then add 100ml of sterile 20% dextrose under sterile conditions and store at 4°C.
4. SD/-trp/raf
  a. Dissolve 1.7g yeast nitrogen base, 5g ammonium sulphate, 0.74g -trp drop out supplement in 850ml MillQ H_2_0.
  b. Adjust pH to 5.8.
  c. Add MillQ H_2_0 to final volume of 900ml.
  d. Autoclave for 20min at 121°C.
  e. Wait to cool, then add 100ml of sterile 20% raffinose under sterile conditions (e.g., under a flame) and store at 4°C.
5. 20% Dextrose
  a. Dissolve 200g dextrose in 900ml MillQ H_2_O. Add NaOH drop by drop if necessary to dissolve all the sugar.
  b. Adjust to pH 6.5.
  c. Add MillQ H_2_O to final volume of 1L and filter sterilize. Store at 4°C.
6. 20% Galactose (see note 10)
  a. Dissolve 200g galactose in 900ml MillQ H_2_O. Add NaOH drop by drop if necessary to dissolve all the sugar.
  b. Adjust to pH 5.8.
  c. Add MillQ H_2_O to final volume of 1L and filter sterilize. Store at room temperature.
7. 20% Raffinose
  a. Dissolve 200g raffinose in 900ml MillQ H_2_O. Add NaOH drop by drop if necessary to dissolve all the sugar.
  b. Adjust to pH 5.8
  c. Add MillQ H_2_O to final volume of 1L and filter sterilize. Store at room temperature.
8. Electroporation buffer
  a. 1M sorbitol
  b. 1mM CaCl2
  c. Autoclave for 20min at 121°C or filter sterilize and store at 4°C.
9. Conditioning buffer
  a. 0.1M Lithium Acetate
  b. 10mM DTT
  c. Make fresh on the morning of electroporation. 1M stock solutions of each LiAc and DTT can be made in advance, filter sterilized, and used to make the conditioning buffer when necessary.

## Author Contributions

Conceptualization, I.C., ML, and R.H.; methodology, I.C., R.H., M.L, L.B.A.; validation, R.H., M.L, L.B.A., A.A.D.L. and A.B.L.; formal analysis, I.C., M.L, L.B.A.; investigation, I.C., R.H., M.L, L.B.A., A.A.D.L. and A.B.L.; resources, I.C.; data curation, M.L., L.B.A., I.C.; writing—original draft preparation, R.H, M.L., L.B.A., I.C.; writing—review and editing, R.H., M.L, L.B.A., A.A.D.L., A.B.L., I.C.; visualization, M.L., I.C.; supervision, I.C.; project administration, I.C.; funding acquisition, I.C. All authors have read and agreed to the published version of the manuscript.

## Data Availability Statement

All DNA sequences used for analysis are available in the Sequence Read Archive (SRA) with BioProject number PRJNA851089.

## Acknowledgments

Authors thank Dr. Tamara Sternlieb for comments on this manuscript.

